# Minimal detection and low biological fluctuation of mitochondrial CpG methylation at the single-molecule level

**DOI:** 10.1101/2020.09.14.296269

**Authors:** Chloe Goldsmith, Jesús Rafael Rodríguez-Aguilera, Ines El-Rifai, Adrien Jarretier, Valérie Hervieu, Victoria Chagoya de Sánchez, Robert Dante, Gabriel Ichim, Hector Hernandez-Vargas

## Abstract

Cytosine DNA methylation in the CpG context (5mCpG) is associated with the transcriptional status of nuclear DNA. Due to technical limitations, it has been less clear if mitochondrial DNA (mtDNA) is methylated and whether 5mCpG has a regulatory role in this context. The main aim of this work was to develop and validate a novel tool for determining methylation of mtDNA and to corroborate its existence across different biological contexts. Using long-read nanopore sequencing we found low levels of CpG methylation (with few exceptions) and little variation across biological processes: differentiation, oxidative stress, and cancer. 5mCpG was overall higher in tissues compared to cell lines, with small additional variation between cell lines of different origin. Although we do show several significant changes in all these conditions, 5mCpG is unlikely to play a major role in defining the transcriptional status of mitochondrial genes.

## Introduction

It has long been established that mitochondria are the powerhouse of our cells. They are responsible for producing ATP through the electron transport chain, contributing to the cellular energetic and redox homeostasis (Porporato et al., 2018). In addition, mitochondria have many other functions including the regulation of apoptotic pathways as well as storing calcium for cell signaling (Porporato et al., 2018). The number of mitochondria in a single cell can vary widely; some cells having no mitochondria, such as red blood cells, while other cells can have hundreds, like liver cells (Alberts et al., 2002).

Mitochondrial DNA (mtDNA) has a molecular weight of 16.5 kb and is comprised of a Heavy Strand (HS) and a Light Strand (LS), with an absence of histones and particular DNA repair requirements (Alexeyev et al., 2013). This unique biology leaves mtDNA exposed to influencing factors from both intra-and extra-cellular origin. For example, reactive oxygen species (ROS) can increase mtDNA copy number (Sun and St John, 2018) and exposure to chemicals can cause mtDNA damage (Weinhouse, 2017). Moreover, alcohol exposure can induce oxidative stress (Lieber, 1991) and increase the expression of mtDNA methyl transferases (mtDNMT1) (Bellizzi et al., 2013). These events highlight the sensitivity of mitochondria to environmental factors which can have downstream consequences for cellular respiration as well as cancer development and progression.

Regulation of mtDNA gene expression occurs primarily through the Displacement loop (D-loop); a 1200-bp non-coding region of the mitochondrial genome. This region controls mitochondrial replication as well as transcription of its encoded genes through a number of different start sites and promoter regions (Crews et al., 1979; Fish et al., 2004).

Among regulatory mechanisms in nuclear DNA, DNA methylation is well characterized and known to be influenced by metabolic activity. In the human genome, cytosine methylation (5mC) occurs mainly in a CpG context (i.e. a cytosine followed by a guanine). However, the existence of mitochondrial cytosine methylation at all has been a topic of debate, with evidence for high levels of mtDNA 5mC in certain human cells and strand-biased non-CpG methylation (Bellizzi et al., 2013; Dou et al., 2019; Feng et al., 2012; Pirola et al., 2013). However, other studies suggested that some of these findings were due to incomplete bisulfite conversion being caused by a failure to linearize mtDNA prior to sequencing (Hong et al., 2013; Mechta et al., 2017; Owa et al., 2018). Moreover, the tools to analyse the presence of DNA methylation rely heavily on sodium bisulfite conversion and PCR amplification; which damage DNA and can lead to bias (Li and Tollefsbol, 2011).

Nanopore sequencing is a unique, scalable technology that enables direct, real-time analysis of long DNA or RNA fragments (Madoui et al., 2015; Seki et al., 2019). It works by monitoring changes to an electrical current as nucleic acids are passed through a protein nanopore. The resulting signal is decoded to provide the specific DNA or RNA sequence. Moreover, this technology allows for the simultaneous detection of nucleotide sequence and DNA and RNA base modifications on native molecules (Jain et al., 2016); hence, removing introduced bias from sodium bisulfite treatment and PCR amplification.

The overall aim of this study was to produce conclusive data on the presence or absence of mtDNA CpG methylation (5mCpG) using a novel technique, and to determine its conservation across different biological conditions. Three cellular settings known to influence mitochondrial dynamics were employed: a model of cellular differentiation, cancer and a model of oxidative stress. After enrichment, mitochondria were sequenced using a ONT Minion device and mtDNA methylation status was directly obtained from the raw signals. We observed low levels of strand specific DNA methylation in hepatocytes with consistent changes related to sample origin.

## Results

### Nanopore sequencing reliably detects CpG DNA methylation (5mCpG)

Long read sequencing is a rapidly evolving field that is largely still in its infancy. Hence, we first sought to determine the reliability of using nanopore sequencing to detect DNA methylation from native DNA in our own hands. To do so, we sequenced genomic DNA extracted from the human liver cell line HepaRG, using an Oxford Nanopore Minion device (ONT). Global patterns of DNA methylation were consistent with the known depletion of 5mCpG at CpG islands (CGIs) (Figure 1A).

**Figure 1.**
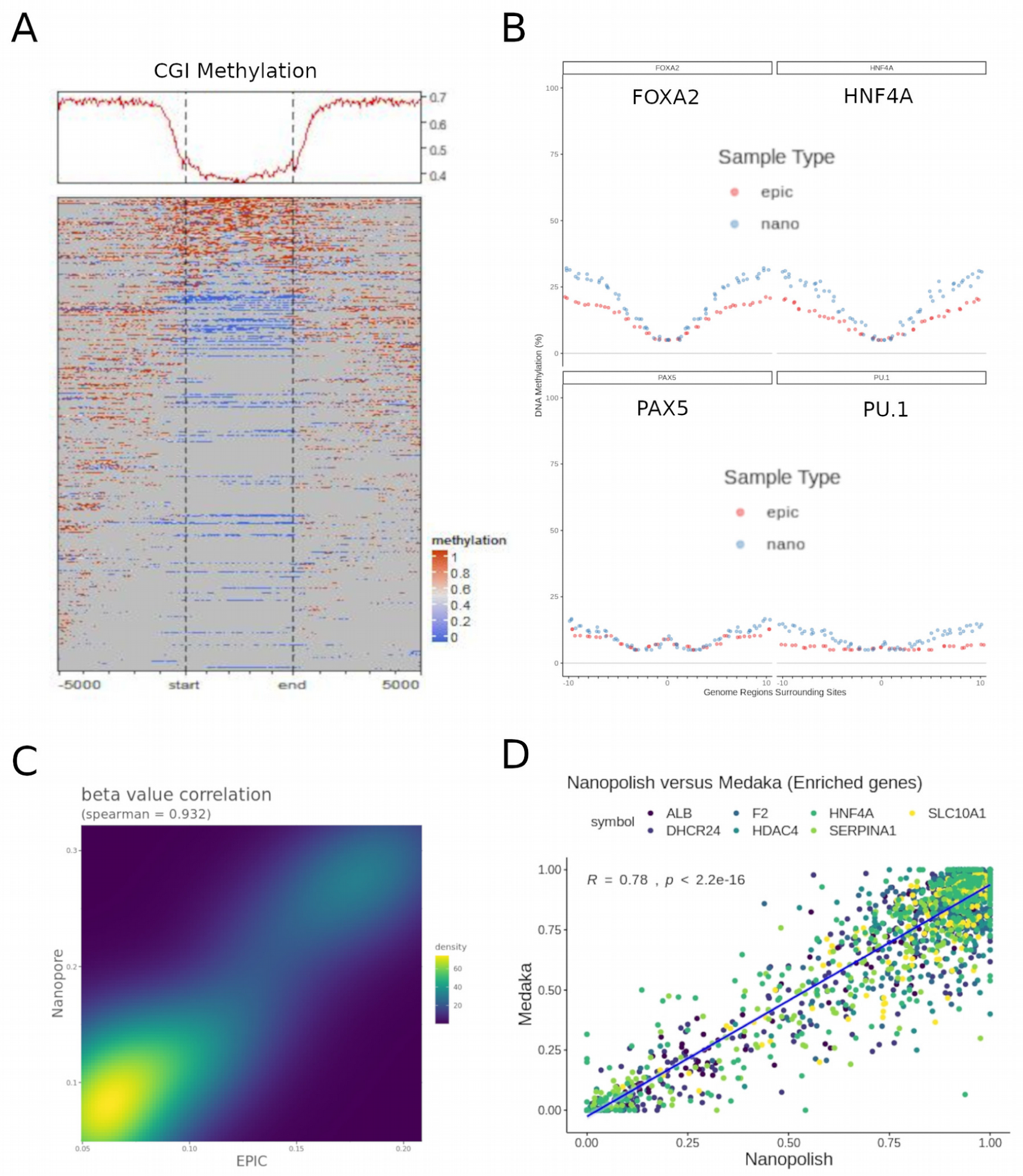
Inspection of 5mCpG data obtained using nanopore sequencing. A) Global profile of nuclear DNA methylation at CpG islands (CGI), obtained after nanopore sequencing of the human liver cell line HepaRG. B) Aggregated DNA methylation data was obtained for hepatocyte active (FOXA2 and HNF4A, top panels) and control (PAX5 and PU.1, bottom panels) transcription factor binding regions. Methylation profiles for EPIC (epic) and Nanopore (nano) (red and blue lines, respectively) are shown for each aggregated dataset. C) EPIC-Nanopore correlation for 5mCpG data on all aggregated datasets shown in (B). D) Nanopore targeted sequencing for a panel of hepatocyte identity genes was used to basecall 5mCpG using two different bioinformatic pipelines: Nanopolish and Guppy+Medaka (see Methods). Single CpG level correlations are shown.

We then compared genome-wide methylation patterns of DNA from HepaRG cells sequenced with nanopore, to those obtained with EPIC Bead Arrays (Illumina) (Rodríguez-Aguilera et al., 2020). To overcome the problem of sparsity in DNA methylation data, we aggregated CpG methylation values from more than 130k transcription binding site loci corresponding to hepatocyte-specific (FOXA2 and HNF4A) and control (PAX5 and PU.1) target regions. Both, EPIC and Nanopore data are able to capture the expected dip in methylation associated with active regulatory regions (Figure 1B, top panels) (Lawson et al., 2018). In contrast, non-active transcription factor binding sites produce a flat methylation profile after aggregation of a similar number of genomic regions in both EPIC and Nanopore data (Figure 1B, bottom panels). Both techniques were highly correlated when aggregated data from all transcription factor binding sites was taken together (Figure 1C).

Next, we tested different available tools to detect DNA methylation from Nanopore sequencing data. We used the well-established tool, Nanopolish (Simpson et al., 2017) which uses a hidden markov model to detect DNA methylation and compared it to the novel tool Guppy + Medaka which has been trained to basecall for modified human CpG dinucleotides using a recurrent neural network (Wick et al., 2019). To perform a site-level correlation, we used targeted nanopore sequencing data from HepaRG cells with a higher coverage in a set of hepatocyte identity genes (i.e. *ALB, F2, HNF4A, SLC10A1, DHCR24, HDAC4* and *SERPINA1*). Using this method of comparison, DNA methylation values were highly correlated, with Guppy + Medaka having a higher tendency towards calling cytosines as unmethylated (Figure 1D). Nanopolish and Medaka outputs have previously been compared with a slightly higher tendency for Medaka to call unmethylated cytosines. (Gilpatrick et al., 2020); as such, these data are in line with former studies.

Therefore, in agreement with recent publications, 5mCpG methylation can be reliably obtained from native DNA using nanopore sequencing and different bioinformatic algorithms. For all analyses presented below we used Guppy + Medaka for extraction of 5mCpG values and Nanopolish for verification and visualization.

### Detection of mtDNA 5mCpG in long reads

Having shown the suitability of nanopore sequencing for analysis of nuclear 5mCpG, we used the same strategy on mtDNA enriched by subcellular fractionation of different cell lines (Figure 2A). Importantly, mtDNA was linearized enzymatically before sequencing using a Minion device. This technique enabled the clear enrichment of the mitochondrial cellular fraction, measured by protein expression of mitochondrial or cytosolic markers (Figure 2B). After sequencing, we obtained a high fraction of reads mapping to mtDNA. Of note, due to long read length, the proportion of mapped reads and their coverage was higher than 80%. Indeed, some reads consisted of full-length mtDNA sequences (Figure 2C). Interestingly, we observed an unequal representation of the heavy strand (HS) and light strand (LS) of mtDNA (Figure 2C).

**Figure 2.**
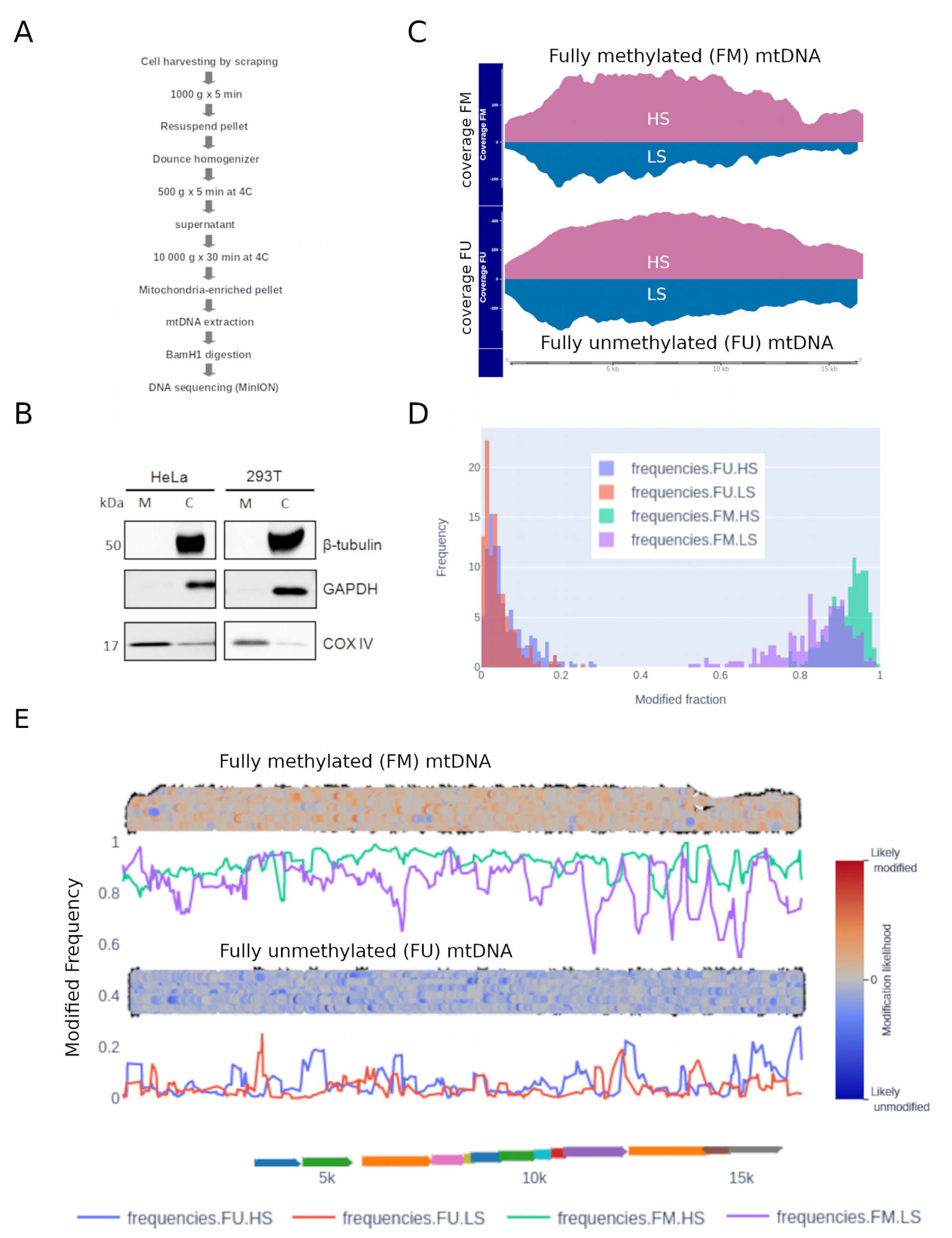
5mCpG methylation in mtDNA. A) Protocol of subcellular fractionation and mtDNA extraction used before nanopore sequencing. B) Quality control of mtDNA enrichment in different cell lines (i.e. HeLa and 293T) using western blot against b-Tubulin, GAPDH, and COX-IV in mitochondrial (M) and cytosolic (C) fractions. C) The same protocol was used on the liver cell line HepaRG. In addition, whole genome amplification was used to produce a “fully” unmethylated control (FU), and followed by DNA methylase (M.Sssl) treatment to produce a “fully” methylated control (FM). Nanopore sequencing coverage for the heavy strand (HS) and the light strand (LS) in FU and FM mtDNA-enriched HepaRG samples. D) Nanopolish was used to infer 5mCpG likelihood and extract methylation frequency tables. Histogram of 5mCpG frequencies in FU and FM, colored by strand. E) Strand-specific 5mCpG frequency plots (colored lines), and 5mCpG likelihood pile-plots (100 reads per sample). Gene mapping to mtDNA are shown in the bottom track as colored arrows.

We attributed this to the efficiency of the Bam1 enzyme in its ability to cut the HS more efficiently and leaving a slightly higher ratio of 5’ ends available for the ligation of adapters before loading onto the Nanopore sequencing device. Moreover, recent findings suggest there can be an unequal representation of mtDNA CpG methylation on the HS and LS (Dou et al., 2019). Hence, to reduce any potential bias to the average methylation of each CpG site, we considered the methylation of the HS and the LS separately for further analysis. Furthermore, mitochondrial populations can be heterogeneous within a single cell. Therefore, we also took advantage of single molecule methylation, by visualizing the methylation of whole mtDNA reads to better understand the single molecule methylation distribution in our samples.

To validate the accuracy of nanopore for detecting 5mC, we prepared fully unmethylated (FU) and fully methylated (FM) mtDNA controls. FU was prepared by whole genome amplification and then FM was prepared by methylation of CpG nucleotides using DNA methyltransferase (M.Sssl). As expected, 5mCpG profiles were opposite in FU and FM mtDNA controls (Figure 2C and 2D). Some residual methylation was observed in the FU control and we considered these levels as a baseline for this technique (Figure 2D and 2E). Indeed, we used the FU control as our background to call detectable methylation. This value was obtained by dividing the number of called sites as methylated by the total number of called sites in the FU sample. The background calculated for the mtDNA HS was 0.022 and for the LS 0.016. We were also able to identify some fully unmethylated reads in the FM control. We attributed this to the efficiency of the DNA methyltransferase (M.Sssl) in methylating these specific reads. Furthermore, this observation highlights the utility of this approach to identify a mixture of DNA in a single sample.

We observed low basal levels of 5mCpG in mtDNA from HepaRG cells. Indeed, we did not identify any differential methylation between the FU control and HepaRG cells, either globally or at the CpG site or strand-specific levels (see next Section).

These data shows that we are able to detect mtDNA methylation with Nanopore sequencing in FM and FU controls. 5mCpG is not different from the unmethylated control at HepaRG basal conditions. We next went further to investigate 5mCpG in several models known to modify mitochondrial activity.

### mtDNA methylation was not affected by hepatocyte differentiation

Hepatocyte differentiation implies metabolic rewiring and changes in mitochondrial content and activity (Yu et al., 2012). As such, this dynamic context may involve concomitant changes in mtDNA methylation. The bipotent liver progenitor cells, HepaRG, are capable of *in vitro* differentiation into hepatocytes and biliary cells. By plating HepaRG cells under differentiating conditions during four weeks we obtain a mixture of the hepatocyte and biliary lineages (Ancey et al., 2017; Cerec et al., 2007; Rodríguez-Aguilera et al., 2020). This well-established model allows us to compare hepatic “progenitor like” cells to their “differentiated” counterpart. We used minimally photo-toxic holo-tomographic microscopy combined with mitochondrial labelling (using MitoTracker Green) to determine mitochondrial content. We observed in both cellular tomogram and MitoTracker staining profile that differentiated HepaRG cells have a higher mitochondrial content, as well as more lipid droplets when compared to their progenitors (Figure 3A).

**Figure 3.**
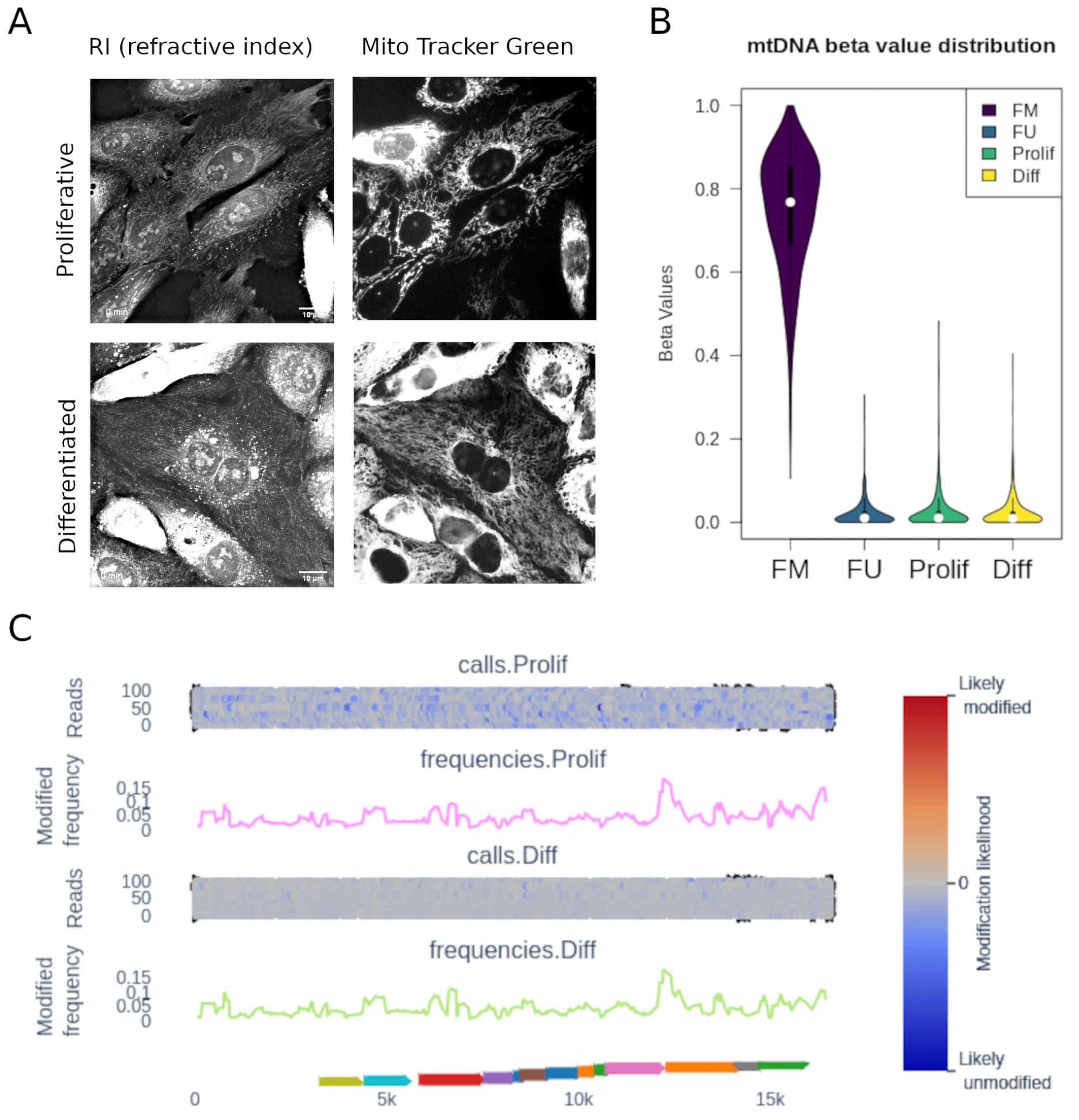
Methylation of mitochondrial DNA measured by nanopore sequencing of a liver progenitor cell line. A) holotomography images of proliferative (progenitor) HepaRG cells and their differentiated progeny. Left panel: Refractive Index (RI) map. Right panel: MitoTracker Green staining to distinguish mitochondrial content and distribution. B) mtDNA enriched DNA extracts from HepaRG cells were linearized and sequenced, as described in Figure 2A. The distribution of 5mCpG beta values (both strands combined) is shown for proliferative (Prolif) and differentiated (Diff) HepaRG, as well as FM and FU controls. C) Methylation frequency and likelihood (pile-plots for the first 100 reads) is shown for proliferative and differentiated HepaRG (one representative sample of three independent differentiation assays). Methylation likelihood scale shown in the pile-plots represents unlikely methylated in blue, likely methylated in red, and intermediate values in gray. Gene mapping to mtDNA are shown in the bottom track as colored arrows.

To determine the effect of hepatocyte differentiation on mtDNA methylation, we used the nanopore sequencing protocol described above, comparing progenitor-like HepaRG cells with their differentiated progeny. In both cases, methylation values were not different from the fully unmethylated control (Figure 3B). There was no differential methylation when directly comparing proliferative and differentiated HepaRG cells (Figure 3C). Similar results were obtained when analyzing both strands together or independently. Interestingly, the likelihood of methylation, as calculated with nanopolish, was higher in differentiated HepaRG cells. This can be seen at the read level (likelihood scale in Figure 3C shows mainly blue reads in proliferative and mainly gray reads in differentiated cells). However, this difference was not high enough to be called as methylation and/or may represent additional nucleotide modifications.

### mtDNA methylation in liver cancer

Most cancer cells display a switch in their metabolic configuration, primarily relying on aerobic glycolysis instead of mitochondrial oxidative phosphorylation (Vander Heiden et al., 2009). In addition, it was recently described that, mtDNA from liver cancer cells had higher levels of CpG methylation than that of non-tumorigenic liver cells *in vitro* (Patil et al., 2019). With this in mind, we wanted to go further and test if mitochondria *in vivo* exhibit this same pattern of DNA methylation. To this end, we sequenced the mitochondria of ten patient liver tissue samples (normal and tumor matched pairs). Both tumor and non-tumor tissues displayed 5mCpG methylation above background levels at several CpG sites (Figure 4A). However, we did not find differentially methylated sites when comparing tumor and their matched adjacent tissues (paired, multifactor approach). A subset of CpG sites with lowest p values for this comparison (non-adjusted p < 0.05), were able to partially discriminate tumors from non-tumor tissues, with the latter displaying slightly higher levels of 5mCpG (Figure 4B).

**Figure 4.**
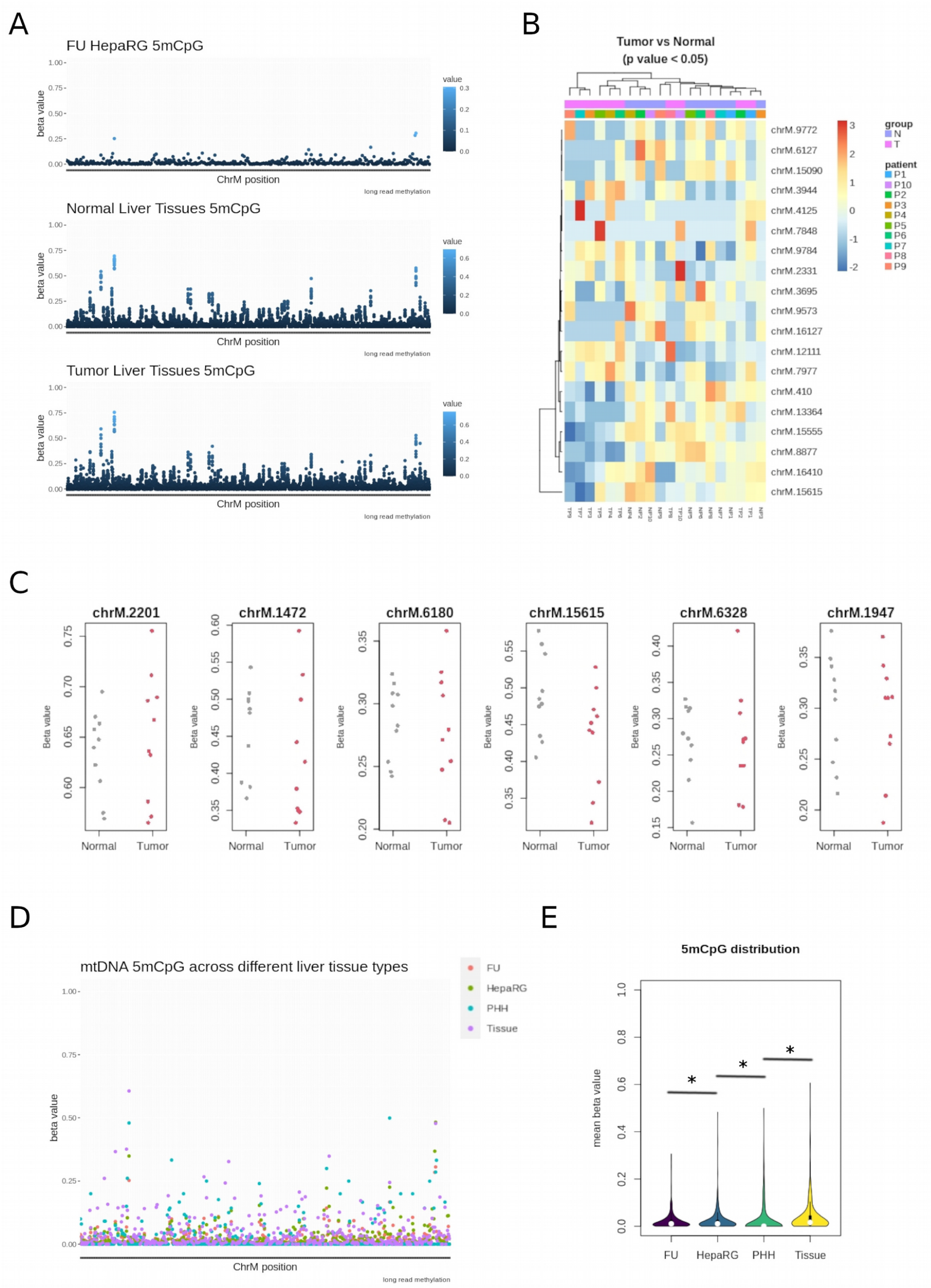
Long read DNA methylation in liver cancer. DNA was extracted from 10 hepatocellular carcinoma (HCC) patients and matched non-tumor adjacent tissues. A) 5mCpG values are shown after nanopore sequencing of a fully unmethylated (FU) liver cell ine (HepaRG, top panel), 10 non-tumor liver tissues (middle panel), and 10 matched tumor tissues (bottom panel). B) 5mCpG heatmap of top most significant (lowest p value in the paired Tumor vs. surrounding comparison) CpG sites. Annotations include Tumor (T) vs Normal (N) status, and patient ID (P1 to P10). C) stripchart of 5mCpG (beta values) for those sites displaying higher levels of methylation in tissues relative to the background (FU sample). Each Normal or Tumor sample is represented in gray and red, respectively. D) 5mCpG values along mtDNA for fully unmethylated control (FU), proliferative HepaRG cells, primary human hepatocytes (PHH) and one representative non-tumor liver tissue (Tissue). E) Distribution of mitochondrial 5mCpG in the same samples represented in (D). (*) indicates p value < 0.05, Mann-Whitney’ test.

Rather than differential methylation between tumors and non-tumor tissues, we found consistent 5mCpG at discrete sites (when compared to the FU background control) in both type of samples (Figure 4C). Most 5mCpG was detected exclusively in the HS (Table), and only 3 CpG sites were consistently found in the LS (i.e. chrM:314, chrM:5469, and chrM:14382). There were also more sites detected as methylated in non-tumor tissues, probably due to a higher 5mCpG variation in tumor samples (Figure 4C and Table 1).

**Table.** Differential methylation in normal and tumor samples, relative to FU control.

| Normal Liver vs. FU control |  |  | HCC vs. FU control |  |
| --- | --- | --- | --- | --- |
| CpG site | pval | fdr | pval | fdr |
| **Heavy Strand** |  |  |  |  |
| chrM:185 | 5.0E-14 | 1.1E-12 | 4.4E-06 | 5.5E-05 |
| **chrM:410** | 1.5E-07 | 1.6E-06 | 3.0E-04 | 2.3E-03 |
| chrM:931 | 2.5E-21 | 1.3E-19 | 2.0E-18 | 1.2E-16 |
| chrM:934 | 1.2E-16 | 3.2E-15 | 1.0E-17 | 5.5E-16 |
| chrM:1176 | 2.0E-28 | 1.5E-26 | 2.8E-26 | 4.1E-24 |
| chrM:1472 | 2.4E-20 | 1.2E-18 | 2.7E-17 | 1.3E-15 |
| chrM:1474 | 2.0E-47 | 8.6E-45 | 8.7E-38 | 3.8E-35 |
| chrM:1483 | 7.6E-09 | 8.9E-08 | - | - |
| chrM:1748 | 2.6E-09 | 3.2E-08 | 1.2E-07 | 2.0E-06 |
| chrM:1947 | 4.4E-07 | 4.2E-06 | 2.8E-06 | 3.7E-05 |
| chrM:2201 | 3.5E-07 | 3.5E-06 | 9.6E-08 | 1.7E-06 |
| chrM:3171 | 2.1E-19 | 7.7E-18 | - | - |
| chrM:3246 | 1.8E-07 | 1.9E-06 | - | - |
| chrM:3965 | 7.8E-12 | 1.4E-10 | 1.8E-07 | 2.8E-06 |
| chrM:5754 | 9.9E-12 | 1.7E-10 | 1.2E-05 | 1.3E-04 |
| chrM:6180 | 3.2E-18 | 1.1E-16 | 6.3E-17 | 2.5E-15 |
| chrM:6328 | 4.7E-11 | 6.8E-10 | 4.7E-09 | 1.1E-07 |
| chrM:6568 | 8.2E-20 | 3.2E-18 | 1.2E-18 | 1.1E-16 |
| chrM:6571 | 5.8E-15 | 1.4E-13 | 5.0E-17 | 2.2E-15 |
| chrM:6850 | 8.9E-32 | 7.7E-30 | - | - |
| chrM:7018 | 1.4E-10 | 1.9E-09 | - | - |
| chrM:7995 | 1.3E-37 | 1.9E-35 | 1.4E-15 | 5.1E-14 |
| chrM:8018 | 1.9E-15 | 4.9E-14 | 1.2E-08 | 2.5E-07 |
| chrM:8116 | 4.4E-17 | 1.3E-15 | - | - |
| chrM:9053 | 6.9E-08 | 7.7E-07 | 5.5E-06 | 6.6E-05 |
| chrM:9161 | 1.9E-10 | 2.5E-09 | 1.0E-08 | 2.2E-07 |
| chrM:9380 | 1.9E-32 | 2.0E-30 | 3.3E-25 | 3.6E-23 |
| chrM:9382 | 6.5E-41 | 1.4E-38 | 7.3E-32 | 1.6E-29 |
| chrM:10201 | 2.7E-10 | 3.4E-09 | 6.4E-10 | 1.6E-08 |
| chrM:11029 | 1.4E-11 | 2.2E-10 | - | - |
| chrM:11161 | 2.3E-05 | 1.5E-04 | - | - |
| chrM:11475 | 1.7E-05 | 1.2E-04 | - | - |
| chrM:11715 | 1.5E-08 | 1.8E-07 | - | - |
| chrM:11912 | 8.1E-07 | 7.4E-06 | - | - |
| chrM:14696 | 5.0E-13 | 1.0E-11 | 6.1E-06 | 7.0E-05 |
| **chrM:15615** | 3.4E-04 | 1.9E-03 | 3.8E-04 | 2.7E-03 |
| chrM:15925 | 4.5E-06 | 3.5E-05 | - | - |
| **Light Strand** |  |  |  |  |
| CpG site | pval | fdr | pval | fdr |
| chrM:314 | 1.5E-18 | 6.4E-16 | 1.3E-07 | 2.8E-05 |
| chrM:4425 | 2.9E-07 | 1.1E-05 | - | - |
| chrM:5469 | 9.1E-07 | 3.0E-0520 | 1.6E-05 | 1.0E-03 |
| chrM:14382 | 6.9E-18 | 1.5E-15 | 5.8E-12 | 2.5E-09 |

While the liver cell line HepaRG did not display 5mCpG above background (Figure 3), liver tissues were consistently methylated at discrete CpG sites regardless of their tumor/normal status. This suggests that 5mCpG may be lost in culture conditions. Indeed, major metabolic alterations, notably metabolic repression, has been described after hepatocytes are placed in culture (Cassim et al., 2017). In line with this, we observed intermediate 5mCpG values in primary human hepatocytes (PHH) after two weeks in culture (Figure 4D). Globally, 5mCpG was not different in HepaRG as compared to the FU control (p = 0.5). In contrast, 5mCpG was higher in PHH relative to HepaRG (p < 2.2e-16), and higher in liver tissues relative to PHH (p < 2.2e-16) (Figure 4E). This result was similar when analyzing separately both mtDNA strands.

Therefore, there are no strong differences in mitochondrial 5mCpG in tumors relative to their matched normal liver tissues. Instead, we were able to detect consistent 5mCpG in tissues and a gradual loss in 5mCpG values as samples are placed in cell culture conditions.

### mtDNA methylation was not affected by oxidative stress

In addition to differentiation and cell transformation, mitochondrial activity is largely associated to oxidative stress, and therefore an interesting process where to study 5mCpG variation. To induce oxidative stress *in vitro*, we used an established method utilizing hydrogen peroxide to induce reactive oxygen species (ROS) (Yagi et al., 2013). Several cell lines were tested (data not shown) and *Homo sapiens* embryonic kidney 293T cells emerged as an ideal candidate for an oxidative stress model. Treatment for two hours was sufficient to induce oxidative stress in 293T cells measured by MitoSox staining, which could be rescued by treatment with N-acetylcysteine (NAC) (Figure 5A and 5B).

**Figure 5.**
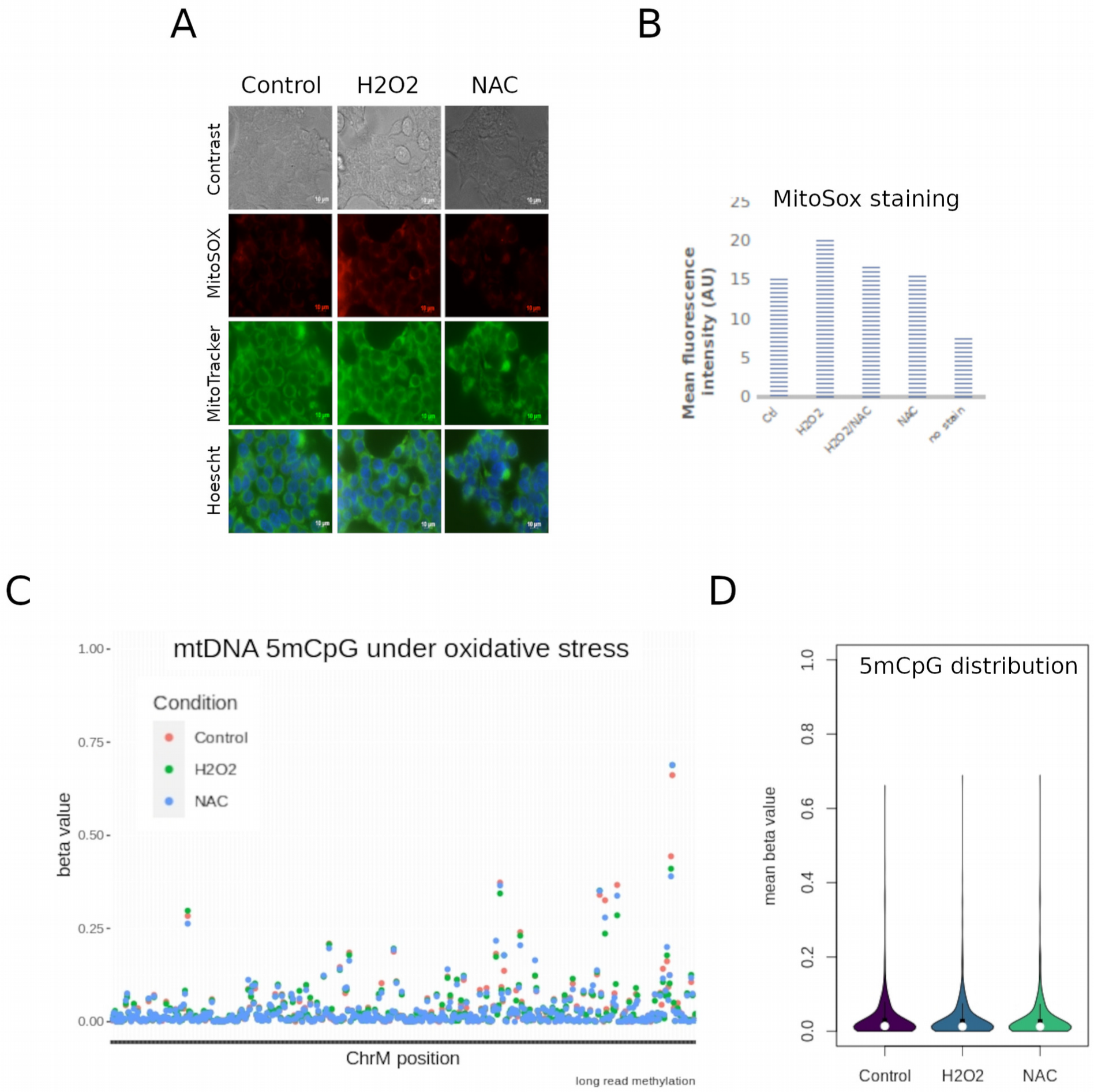
Mitochondrial 5mCpG in response to oxidative stress. Mitochondrial DNA was obtained from the human embryonic kidney cell line 293T under basal conditions (Control), oxidative stress (H2O2) and oxidative stress rescued with N-acetlycysteine (NAC). A) Representative images of phase contrast, Mitosox, MiitoTracker and Hoescht staining. B) MitoSox quantification of 5 independent replicates. C) 5mCpG measured by nanopore under the same experimental conditions. Each dot represents the average of triplicate values for each condition and each CpG site. D) mitochondrial 5mCpG distribution (both, HS and LS strands together) of the data shown in (C).

At basal levels, 293T cells exhibited higher global levels of 5mCpG than HepaRG cells (p value = 0.05). However, the same strand specific methylation was observed (i.e. higher 5mCpG in the HS). We then aimed to study the effect of oxidative stress on mtDNA methylation levels. In order to do so, we induced oxidative stress in 293T cells and compared the mtDNA methylation levels before and after treatment. We also rescued these cells from the induced oxidation using NAC. Again we calculated the differential methylation between these three treatment groups and with a threshold of 10% (Figure 4), we did not observe any differential methylation on the HS or the LS (Figures 5C and 5D).

Although 5mCpG was clearly detectable in 293T cells, and higher than HepaRG cells, it was not significantly affected by H2O2 exposure.

## Discussion

In the present study we have shown that nanopore sequencing can reliably detect 5mCpG in mitochondrial DNA from human cells. Exploiting the advantages of long reads and native DNA sequencing, we show that 5mCpG can be detected at discrete CpG locations at levels that depend on the cellular model (i.e. immortalized cell line, primary cells, or tissue). However, we did not observe differential 5mCpG in three biological contexts: in vitro differentiation of a liver progenitor cell line, comparison of human liver tumors and their matched non-tumor tissues, and in vitro induction of oxidative stress.

For the first time, we provide a comprehensive characterization of mtDNA methylation lin liver cells, kidney cells and liver tissue by long read sequencing (ONT). We were able to use this highly novel tool to detect the methylation patterns along 16kb reads spanning the entire mitochondrial chromosome with deep coverage of >10000x on naive DNA. In doing so, we have produced a map of mtDNA 5mCpG, that has completely eliminated any introduced bias from bisulfite conversion and PCR amplification; tools that we have relied on, and that have served us well for many years.

Using ONT, we identified low basal levels of mtDNA methylation at specific regions in liver cells *in vitro*. These levels were lower than that previously described (Ghosh et al., 2014; Patil et al., 2019), however these authors have noted that CpG methylation was highly cell specific. While we also analyzed the mitochondrial methylome of liver cells, we did not use the same cell lines as these previous studies. Hence, the differences observed are likely due to the cell specific nature of mtDNA 5mCpG. Moreover, it should be noted that a series of work outlining amendments to the bisulfite conversion protocol for mitochondria have been published in order to ensure the bisulfite conversion efficiency is properly controlled for (Owa et al., 2018). Since the average mtDNA CpG methylation levels were very low, we further validated this work through extensive comparisons of basal 5mCpG with negative controls. Other studies have reported similar cell specific mtDNA methylation patterns (Bellizzi et al., 2013), however, we are the first to represent these landscapes in differentiation models and/or using long reads.

Furthermore, we have also for the first time, clearly identified strand specific mtDNA methylation using long read sequencing. We observed higher levels of 5mCpG in the HS generally. This was in accordance with recent reports that have also identified a strand specific methylation via different techniques such as bisulfite sequencing and meDIP (Dou et al., 2019; Ghosh et al., 2014; Patil et al., 2019).

We did not find significant variation in 5mCpG under oxidative stress conditions. While the effect of oxidative stress on mitochondrial activity has been extensively studied (Ashari et al., 2020; Yu et al., 2020), there had not yet been a comprehensive mapping of mtDNA 5mCpG in oxidative stress conditions. In fact, we could not find any work that has investigated mtDNA methylation in this context.

Despite our novelty, there are limitations to this work as it stands; including the lack of investigation into non-CpG methylation, which has previously been characterized in liver cancer cells and linked strongly to the control of mitochondrial gene expression (Bellizzi et al., 2013; Patil et al., 2019), as well as the detection of 5hmCpG, which has been also reported in mtDNA (Shock et al., 2011), and is highly dynamic in liver cell differentiation and linked to gene regulation (Ancey et al., 2017; Rodríguez-Aguilera et al., 2020). Therefore, it is clear that more work is needed to develop long read sequencing tools to determine non-CpG methylation or other modified bases like 5hmC in general. There are technical and bioinformatic limitations to nanopore sequencing currently. But this field is rapidly advancing and as such we are confident that making this data publicly available will continue to contribute to this important work.

In conclusion, nanopore is a useful tool for the detection of modified DNA bases on mitochondria, however, care must be taken to consider the HS and LS strands separately as well as the heterogeneity of mitochondrial populations.

## Methods

### Cell culture, maintenance and differentiation

HepaRG cells were cultured in Williams media enriched with 10% Fetal calf serum clone II, 1% Penicillin/Streptomycin, L-glutamine (2mM), insulin (5µg/mL) and hydrocortisone (25µg/mL). Proliferative HepaRGs were taken before reaching 50% confluence and differentiated hepaRGs were differentiated as previously described (Ancey et al., 2017; Cerec et al., 2007; Rodríguez-Aguilera et al., 2020).

HEK293T, immortal cells derived from embryonic kidney were grown in tissue culture dishes (Falcon, Becton Dickinson) and cultured in DMEM 1X media containing 1% Penicillin/Streptomycin, 1% sodium pyruvate, 1% L-glutamine, 1% non-essential amino-acids, all from Life Technologies, and 10% fetal bovine serum (Eurobio Abcys).

### Induction of oxidative stress

HEK293T cells were treated ydrogen peroxide (H2O2) (Sigma-Aldrich, 216763) at a concentration of 500 μM for 2 hours, alone or in combination with 5mM N-acetyl-cyteine (NAC) (Sigma-Aldrich, M for 2 hours, alone or in combination with 5mM N-acetyl-cyteine (NAC) (Sigma-Aldrich, A7250). When using NAC, cells were pre-treated for 2hrs with 5mM NAC.

The mitochondrial superoxide indicator stain MitoSOX (ThermoFisher, M36008) was used to probe the relative oxidative stress in live cells. Cells were stained witiih 1uM MitoSox diluted in DMEM. 250,000 cells were incubated with 330 ul for 30 min and analyzed by flow cytometry, then washed with PBS and trypsinized. Flow cytometry tubes were kept on ice and in the dark until use. Flow cytometry analysis was performed with a FACSCalibur (BD Biosciences). The mean fluorescence intensity of minimum 10,000 stained cells and unstained control cells were recorded and plotted for analysis. Alternatively, MitoSOX was analyzed by epifluorescence microscopy (Zeiss, Axio Observer).

### Holotomography

Differentiated and proliferative HepaRG cells were plated at high confluence. Mitotracker (100nM) was added to normal growth medium for 1h before imaging with a 3D Cell-Explorer Fluo (Nanolive, Ecublens, Switzerland) using a 60x air objective. Refractory index maps were generated and images were processed every 5 seconds for 20 minutes with the STEVE software.

### Subcellular fractionation and mtDNA extraction

Subcellular fractionation was performed as previously described (Arnoult et al., 2003) with some modifications. Briefly, cells were washed with PBS, harvested by scraping and centrifuged at 1000 g for 5 min. The pellet was re-suspended in buffer containing 210 mM sorbitol, 70 mM sucrose, 1 mM EDTA, 10 mM HEPES and 0.1% BSA (Sigma) before grinding with a Dounce Homogenizer (Wheaton, USA) with a loose and tight pestle (100 strokes with each pestle). Cells were observed under microscope (Axiovert 40C, Zeiss) with trypan blue dye to assess cell membrane disruption followed by centrifugation at 500 g for 5 min at 4 °C. The supernatant was collected before centrifugation at 10 000 g for 30 min at 4 °C. DNA extraction (Nucleospin Tissue, Macherey-Nagel) was performed on the resulting pellet according to manufacturer instructions. mtDNA was digested using BamH1 HF (New England BioLabs) in order to linearize mtDNA genome.

### Fully unmethylated and fully methylated controls

After mtDNA enrichment and linearization, we prepared a negative (FU = fully unmethylated) control sample from differentiated HepaRG mtDNA by performing whole genome amplification using a repliG kit (Qiagen) according to manufacturer’s instructions. After amplification, a positive control for methylation (FM = fully methylated) was prepared. Briefly, CpG dinucleotides were methylated by incubating 1µg of DNA with S-Adenosyl methionine (SAM) (32µM) with CpG Methyltransferase (M.SssI) (4-25 units) (New England BioLabs) at 37°C for 1h before heating to 65°C for 20mins.

### Patient tissue samples

Human biological samples and associated data were obtained from “Tissu-Tumorothèque Est” (CRB-HCL, Hospices Civils de Lyon Biobank, BB-0033-00046). DNA extracted with the epicentre kit.

### Nanopore sequencing

400ng of DNA from each sample or control was barcoded and multiplexed using the Nanopore Rapid Barcoding Sequencing kit (SQK-RBK004) according to manufacturer’s instructions. Sequencing was conducted with a Minion sequencer on ONT 1D flow cells (FLO-min106) with protein pore R9.4 1D chemistry for 48h. Reads were basecalled with GUPPY (version 4.3.2). Basecalled reads were mapped using Minimap2 to the GRCh38/hg38 human genome.

### Bioinformatic analyses

Basecalling was performed with Guppy version 4.0.15 (ONT). We first determined the methylation status of each CpG site on every read by using the widely used tool, *nanopolish* (Simpson et al., 2017) used recently by (Gigante et al., 2019). For validation, we also called DNA methylation using novel tool, Medaka (git repository reference). Medaka is a tool to create a consensus sequence from nanopore sequencing data. This task is performed using neural networks applied from a pileup of individual sequencing reads against a draft assembly. It outperforms graph-based methods operating on basecalled data, and can be competitive with state-of-the-art signal-based methods, whilst being much faster.

PycoQC was used for data inspection and quality control (https://github.com/a-slide/pycoQC), and methplotlib (https://github.com/wdecoster/methplotlib) for read-level visualizations.

Called CpG sites in the FU control were used to determine a baseline of methylation. The following calculation was utilised: FalsePositiveRate=[#called methylated cytosines in FU/#called cytosines in FU].

For differential methylation analyses we used DSS (Dispersion shrinkage for sequencing data) (Park and Wu, 2016) adapted for nanopore sequencing (Gigante et al., 2019). The aggregated β methylation values for each CpG group are tested for differential methylation using the DSS software (Park and Wu, 2016) and adapted for nanopore sequencing according to (Gigante et al., 2019). Briefly, DSS tests for differential methylation at single CpG-sites, using a Wald test on the co-efficients of a beta-binomial regression of count data with an ‘arcsine’ link function. In order to set minimum requirements for DSS analysis, an internal comparison of biological replicates of differentiated HepaRG cells was undertaken. From this we were able to better understand the background and determine the minimum smoothing and delta values. These values were set at a smoothing of 10-50bp and a delta of 0.05 with minimum P-value of 0.05.

We used the bioconductor packages MIRA (Lawson et al., 2018) for methylation data aggregation, and LOLA for dataset selection (Sheffield and Bock, 2016).

Mann-Whitney’s test was used for pairwise comparisons of 5mCpG distribution.

## Availability of data and material

Datasets generated during the current study will be uploaded to the GEO repository.

## Acknowledgements

The authors would like to thank the patients that participated in this study.

## Authors’ contributions

C.G. carried out the experiments and wrote the first draft of the manuscript; J.R.R.A. and I.E-R. performed experiments and additional validations; C.G., A.J. and H.H.-V. performed all statistical and bioinformatic analyses; V.H. obtained the human samples; R.D., G.I., and V.C.d.S. provided conceptual assistance and supervised experiments; H.H.-V. and G.I. conceived the study; H.H.-V. coordinated the project and wrote the manuscript. All authors discussed the results and manuscript text.

